# A generalised protein identification method for novel and diverse sequencing technologies

**DOI:** 10.1101/2024.02.29.582769

**Authors:** Bikash Kumar Bhandari, Nick Goldman

**Author notes:** Corresponding authors: Bikash Kumar Bhandari, Nick Goldman.

## Abstract

Protein sequencing is a rapidly evolving field with much progress towards the realisation of a new generation of protein sequencers. The early devices, however, may not be able to reliably discriminate all 20 amino acids, resulting in a partial, noisy and possibly error-prone signature of a protein. Rather than achieving *de novo* sequencing, these devices may aim to identify target proteins by comparing such signatures to databases of known proteins. However, there are no broadly applicable methods for this identification problem. Here, we devise a hidden Markov model method to study the generalized problem of protein identification from noisy signature data. Using a hypothetical sequencing device that can simulate several novel devices, we show that on the human protein database (N=20,181) our method has a good performance under many different operating conditions such as various levels of signal resolvability, different numbers of discriminated amino acids, sequence fragments and insertion and deletion error rates. Our results demonstrate the possibility of protein identification with high accuracy on many early experimental devices. We anticipate our method to be applicable for a wide range of protein sequencing devices in the future.

## 1 Introduction

There have been significant advances in nucleotide sequencing technologies in the past decade [1, 2, 3, 4, 5]. Nanotechnological techniques are getting closer to single-molecule resolution and high readout accuracy [6, 7]. In particular, current nanopore sequencing methods for nucleotides rely on electrical signals from small *k*-mers (e.g. *k* ≤ 7), which results in 4*^k^* variations of the superimposed signals [8, 5]. For small *k*-mers, the decoding problem is tractable [5]. In parallel, there has also been exciting progress in the protein sequencing space such as prototype devices and experiments for single-molecule sensing [9, 10] and the development of nanopores suitable for protein translocation [11, 12]. However, compared to DNA sequencing, the progress in protein sequencing has lagged [13, 14, 15]. This is not surprising because protein sequencing involves the discrimination of 20 different residues with complex structures and charge distributions, compared to just four nucleotides in nucleotide sequencing [16, 17]. The possible variations in the signal from *k*-mers in the case of nanopore sequencing of proteins quickly becomes huge (20*^k^* variations), making decoding difficult.

Mass spectroscopy is a widely used method for protein identification which compares the mass spectra of the unknown protein to a database of known spectra of proteins. For *de novo* sequencing, Edman degradation has been largely replaced by tandem mass spectroscopy [18]. However, much proteomics is done by protein identification rather than *de novo* sequencing because in many cases, partial sequence information is enough to identify a sequence [14]. Despite being the gold standard of proteomics, the limited dynamic range of mass spectroscopy makes it unusable to identify very low-concentration peptides [16]. Therefore, efforts have been made to develop alternate methods with higher sensitivity. Methods employing fluorescent tagging of a subset of AAs to generate a fingerprint [19, 20, 21] or using other properties of AA in addition such as binding kinematics [10] are being investigated. Recent works suggest that up to six AAs could be labelled without spectral overlapping, which may be enough to identify the sequence from a database [22, 16].

Nanopore methods seem more promising because of their potentially higher dynamic range and the possibility of long-read sequencing [16]. Progress towards this goal includes the development of engineered biological nanopores, steady translocation of peptides [23, 24, 12, 25, 26], and acquisition of electrical signals from the translocated peptide to discriminate 13 of the 20 AAs [27]. Approaches based on solid-state nanopores are also being explored [28, 29, 30]. Further improvements in single-molecule sensing, for example, through optical signals from plasmonic resonance and surface-enhanced Raman spectroscopy are promising in discriminating all 20 AAs [31, 32, 29, 17, 33]. These advances, and possibly newer engineering techniques and experimental protocols with single AA-level resolution, are required for successful *de novo* protein sequencing [19, 34]. However, improvements in current methods may be enough to develop a sequencing machine that can generate partial signals which can be used to identify proteins from a known database.

Despite these promising steps towards a successful protein sequencing device, considerations such as fluorescent overlap among the labelled AAs, sample preparation, cost, device engineering, ease of operation, nanophysics, nanophotonics, and nanotribological principles mean that early devices will likely employ several strategies to make protein identification easier, while still providing a base for the future *de novo* devices. Well-known strategies include discrimination of a subset of AAs and using protein fragments [19, 20, 21]. In addition, these devices will potentially generate error-prone readings and the signal from these devices may not be fully resolvable for an exact identification of the AAs. Therefore, the decoding algorithms to identify the AAs and proteins will likely output a probabilistic readout over all AAs for each segment of the signal.

One of the probabilistic approaches may be to start with a prior probability distribution of each AA. The prior distribution could be uniform (1/20 for all AA), to reflect the absence of any prior information about the AA distribution on the proteins being sequenced, or adjusted based on some prior knowledge. For example, if it is known that the query proteins or fragments are cysteine-rich, the prior probability for cysteine can be set higher. Upon receiving the signals from the sequencing device, the decoding algorithm would update the priors to give the posterior probability distribution of the AAs such that the correct AA, hopefully, will now have the maximum posterior. In particular, the posterior probability for the correct AA should be close to 1 for a well-resolved signal. Depending on the noise and decodability of the signal, however, they will generally be less than 1 for the correct AA and greater than 0 for one or more incorrect AAs. For the positions where the signal is unavailable or unresolved, the posteriors will be the same as priors.

In general, for a peptide of length *L* (and assuming, for now, the absence of insertion and deletion errors) the decoded output from these sequencing techniques can be written as a 20 × *L* probability matrix, where each column *i* contains the posterior probabilities assigned to each of the 20 AAs possible at position *i* in the sequence. Due to the error-prone nature of the signal acquisition and decoding, a direct reconstruction of each position of the protein sequence using just its posterior probabilities would likely indicate a non-existent or incorrect protein. Therefore, these posterior readouts must be queried jointly against some protein database to identify the protein that they were derived from.

Despite the diverse working principles of many such imaginable devices, they can be reduced to a single hypothetical device which can produce the posterior probability readouts corresponding to the different strategies. Additionally, the hypothetical device is also generalisable to devices without probabilistic decoding algorithms such as based on just the fluorescent tagging, fingerprinting and possibly Edman degradation. In this case, observing one particular wavelength might suffice to reliably identify an AA, which can be represented by a probability vector such that the probability of the correct AA is close to 1.

Therefore, to study the problem of protein identification, we propose such a hypothetical protein sequencing device and couple it with a database search method using its probabilistic readouts. We address the problem of identification of human proteins, using c. 20,000 distinct canonical human proteins from UniProt [35] to simulate reads. We study several operating conditions of the device, such as different levels of signal resolution leading to posterior probabilities between unresolved and well resolved, different length of the input proteins or fragments, different error rates, and various subsets of perfectly detectable AAs, using the readouts from the device to try to identify the correct originating protein. Hidden Markov models (HMMs) and more recently different neural network architectures have been successfully used to sequence DNA [36, 37, 38, 8, 39, 40, 41, 42, 43]. These methods can also be extended to protein sequencing. Despite their potential for higher accuracy, neural network models are often difficult to interpret in a meaningful way. On the contrary, HMMs are simpler probabilistic models that score a sequence based on the matches, insertions, and deletions required to match the probabilistic readouts to a sequence [44]. In this work, we use an HMM for protein identification, first showing that probabilistic, full-length and error-free readouts can be used to identify almost all proteins. Then we show that the reduction of readout length to as short as 10 AAs can be enough to identify more than 95% of the sequences from a single fragment. We further evaluate our protein identification method with simulated fluorescent tagging and show that readouts from just two AAs, leucine (L) and serine (S), on a full-length protein, lead to the identification of 95% of the proteins. In the same case, using one fragment with 100 AAs instead is sufficient to identify at least 76% of the proteins, and if the readout from a third AA, glutamate (E), is added, we can recover at least 94% of the proteins from such shorter fragments. Furthermore, we also test our method on error-prone readings based on discrimination of all 20 AAs or on reduced sets of AAs and show that it is quite robust to a wide range of errors. Our study highlights the potential for success under many different strategies, while also providing useful insights for designing a next-generation single-molecule protein sequencer.

## 2 MATERIALS AND METHODS

### 2.1 Data

We retrieved all reviewed canonical human protein sequences from the UniProt database (UniProt release 2022 04) [35]. To reduce the computation time during our many sequence identification experiments, we discarded proteins above the 99th percentile of the length distribution. This resulted in the exclusion of extremely long sequences such as titin (34,350 residues). We were left with 20,181 sequences with a median length of 411 residues (Supplementary Fig. S1A). These sequences were used as the database to study the problem of identification of human proteins.

### 2.2 Hypothetical single-molecule sequencer

For any input sequence, our hypothetical device sequentially reads each AA and outputs a posterior probability vector over 20 AAs (Figure 1). This posterior takes the value *p_max_* for the correct AA. The parameter *p_max_* gives us the control to simulate performance ranging from perfect determination of every AA (*p_max_* = 1), through devices with good discrimination (e.g. *p_max_* = 0.9), as far as *p_max_* = 0.05, when all AAs are completely indistinguishable. In practice, for a given sequence, the signal quality could vary along the sequence, leading to different *p_max_* along the sequence. The remaining probability (1 − *p_max_*) might also have some non-uniform distribution over the remaining 19 AAs. However, we simplified our simulations by eliminating these degrees of freedom by assuming *p_max_* is a constant in each simulation and (1 − *p_max_*) is equally divided among the 19 incorrect AAs. If the device is free from sequencing errors such as insertions and deletions (indels), the decoded readouts from a protein of length *L* is a 20 × *L* posterior probability matrix.

**Figure 1:**
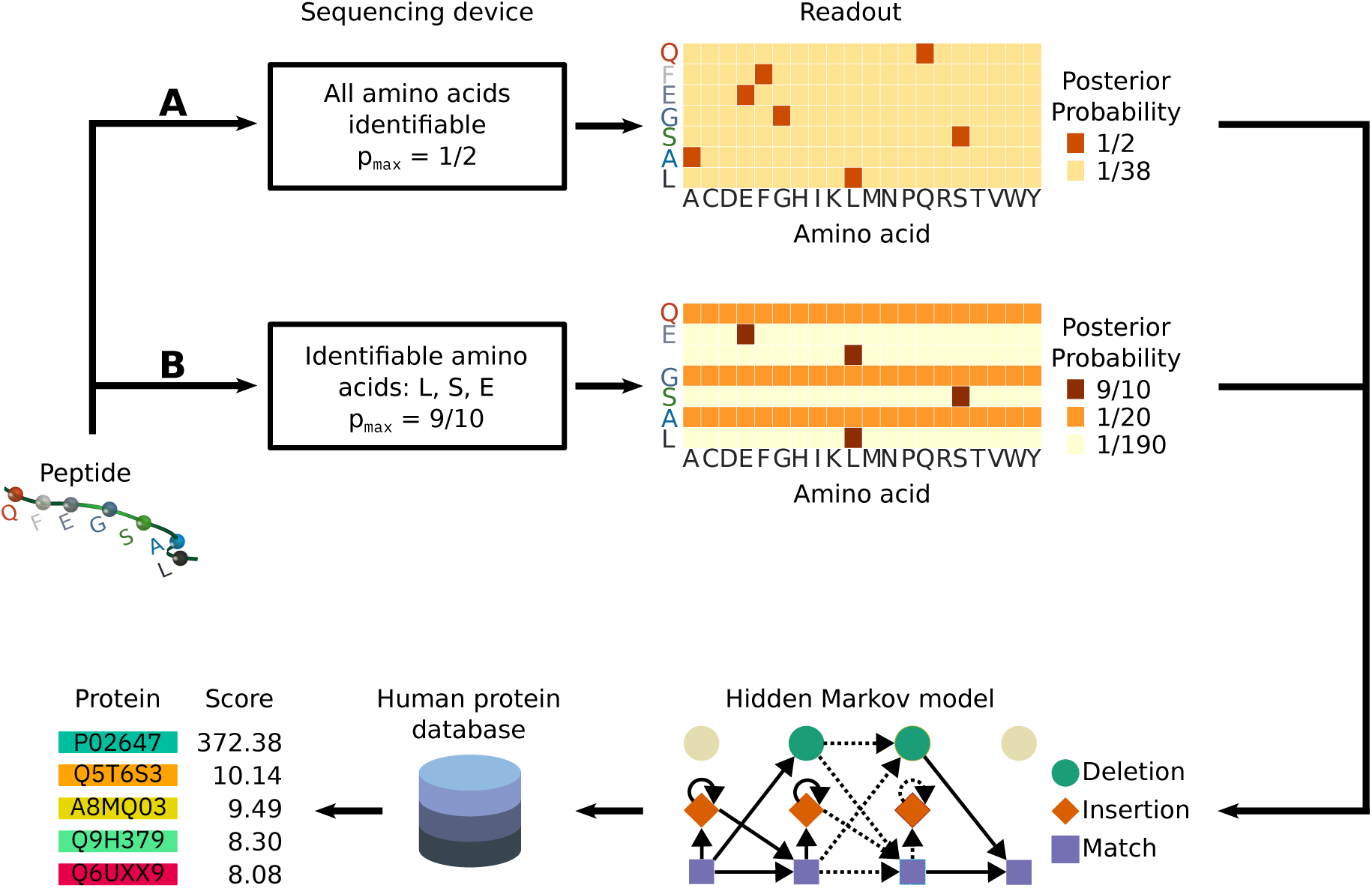
Hypothetical sequencing device used in this study, and data analysis workflow. The sequencing device reads an input protein (in this example, a peptide QFEGSAL) and outputs a posterior probability over all 20 AAs for possibly each AA in the input sequence. Two example operating conditions are shown. In case **A** the sequencing device can identify all AAs with *p_max_*= 1*/*2 and the corresponding readouts for the input peptide are shown as a heatmap of the posterior probabilities. In case **B** the sequencer is set to output high posterior probabilities (*p_max_* = 9*/*10) to the AA subset L, S, and E which could be because either these AAs are tagged or the device is able to generate good quality signal for only these AAs. Hypothetically, the device is configurable to extend this subset from none to all 20 AAs. If the input AA is not in this subset, the posterior probability is uniform (1/20). In this readout, deletion error occurred after the first AA, Q. Therefore the uniform posteriors for F are absent. Similarly, an insertion error occurred after the next AA, E, leading to a spurious probability vector suggesting presence of L. The rates of these insertion and deletion errors are configurable in our simulations. These posterior probabilities are used as emission probabilities of the match state of an HMM. The HMM scores every sequence in the human protein database (N=20,181). For illustration, five top example hits and their corresponding scores from the HMM using posteriors from **B** are shown. The sequence with the highest score (in this example, protein P02647 with score 372.38) is the inferred protein from the posterior probability readout of the input sequence.

### 2.3 Inputs to the sequencer

Developments in nanopore methods have led to successful translocation of full-length proteins, for example, mediated by the enzyme unfoldase ClpX [45] and more recently enzyme-free electroosmotic methods [26]. These experiments suggest the possibility of sequencing devices which take full-length sequences. We therefore started investigating our protein identification method with single full-length sequences as inputs to the sequencer.

In some of the protein sequencing and identification techniques, using a protein fragment is desirable. For example, in fluorescent labelling and subsequent Edman degradation, peptides of length around 30 AAs are preferred due to the limitations of the complete cleavage cycles [46]. Similarly, for nanopore-based devices, some AAs may clog the pore due to their volume, hydrophobicity and charge [47]. Using protein fragments may be one possible strategy to mitigate this problem. Therefore, in addition to using full-length sequences, we also used sequence fragments of various lengths (100, 50, 25, 15, 10, and 5 AAs) as input to the sequencer. First, we again used a single random fragment per protein. However, as many proteins contain conserved domains in common, the location of a single random sequence fragment per sequence might introduce bias in the protein identification process. To investigate and possibly eliminate such systematic biases, while limiting the computational time, we repeated this analysis 10 times per protein, each time using a different random fragment.

Second, since a digested sample could contain many fragments from the same protein, multiple readouts could be used to improve protein identification. Therefore, in cases where performance of our method was not strong, we investigated combining the results from each of the 10 fragment readouts for possible improvements in protein identification.

### 2.4 Amino acids identified by the sequencer

Recent experiments have demonstrated the possibility of generating unique signals from all 20 AAs [31]. This might lead to the development of a device which can discriminate all AAs on a full-length sequence in the near future. Hence, we simulated the case where the device can attempt the identification of all 20 AAs on full-length sequence. Additionally, using our parameter *p_max_*for the posterior of the correct AA at each sequence position, we simulated different working conditions from well-resolved signal (*p_max_* = 0.9) to unresolved signal (*p_max_* = 0.05).

Despite the promising early experimental results in single AA identification, prototype devices are likely to reliably identify only a reduced set of AA. Additionally, fluorescence methods generally label only a small set of AAs [20, 22]. To analyse such scenarios, we also simulated the identification of reduced sets of AAs. Generally, the choice of the set of AAs will be dependent on experimental factors such as ease of labelling and spectral overlap. In our simulations, we used three such sets. The first set consists of the five most abundant AAs (L, S, E, A, and G) in the human protein database (Supplementary Fig. S1B). In contrast, our second set consists of the five least abundant AA (W, M, C, H, and Y), and thus is not very informative and likely represents a worst case. With both of these sets, we varied the number of AAs distinguished from one to all five. In addition, we also used a third set containing AAs (C, Y, K) that are convenient to chemically label [22, 13]. For this set, we considered all single and double AA combinations. In these simulations, the combinatorial space becomes huge. Therefore, we limited ourselves to the full length, 100- and 50 AA-fragments, and *p_max_* of 0.8 and 0.2, which is adequate to study our method’s protein identification from better and modest signal quality.

### 2.5 Incorporating errors

Any sequencing method is prone to errors. For devices based on fluorescence techniques, these errors might arise due to inefficient labelling and detection [16, 48]. For devices dependent upon the translocation of a protein, errors may arise due to the non-uniform speed of the sequence and possibly during the decoding of the signals. In nanopore sequencing of DNA, such errors are also called skip (due to fast translocation) and stay (due to slow translocation) errors [38, 8, 49]. Therefore, it is likely that protein sequencing techniques may also suffer from these types of errors. To model these errors, we simulated different levels of insertions and deletions in the signal. For example, for a sequence of length 100AA at an insertion rate of 10%, we insert exactly 10 random posterior probability vectors. This simulates the misidentification of AAs and possibly a slow translocation of the peptide. Similarly, for a deletion rate of 10%, we delete exactly 10% of the posterior probability vectors. This simulates a faster translocation or a missed detection of signals.

Translocation speed is likely dependent on the shape, size, charge distribution and other physical properties of the AAs and their response to the translocation mechanism used. Therefore, some positions of the protein may be more prone to errors. However, to simplify our analysis, we assumed these errors to be uniformly distributed along the sequence. In cases where a sequence has both insertion and deletion errors, we simulated deletion first followed by insertions. This ensures the insertion errors do not undergo subsequent deletion. Furthermore, after doing an experiment, the location of these errors may not be easily detectable, although, theoretically speaking, it might be possible to infer the position of the errors through analysis of the raw signal and some measure of translocation time. While such additional information might be useful in signal decoding, in our study we assume this information is unavailable: instead of raw signals, our device already gives the decoded readouts as posterior probabilities. We therefore discard all positional information regarding the errors in our simulations.

### 2.6 Identification of proteins using decoded readouts

The readouts from our device are the posterior probabilities of AAs along the sequence reflecting the uncertainties of AAs identified from the signal (Figure 1). The core of our protein identification method is an analysis that uses these probabilities to infer the originating protein. For a single readout from a protein or a protein fragment, if we consider the AAs from the unknown protein as match states of an HMM, the posterior probabilities can be treated as the emission probabilities of these states. Additionally, since there might be sequencing errors in the form of indels, we can also assign some probability of transitioning from any of the match states to the insertion and deletion states. Thus, the readouts of the sequencing device can be described using a classic HMM (see, e.g., [50]). Our goal of identifying a protein sequence from a database using the readouts then becomes very similar to the problem of searching a database of target HMMs using a query sequence. This is a well-studied problem in bioinformatics, HMMER [51] being one of the most widely used tools for this purpose. In our case, we have one HMM query from each readout of a protein, which has to be queried against a database of known target sequences. Therefore, to identify the protein, we can reverse the classical HMM search problem. Using HMMER and the HMM constructed from the readouts, we score all the sequences in the database. We regard the hit with the highest score as the inferred protein.

In the classical HMM search problem, the probabilities of emission from the match state and various transition probabilities of each target HMM are typically calculated from the occurrences of AAs in a multiple sequence alignment of evolutionarily related sequences. However, the HMMs in our study use posteriors from the sequencing device as the match emission probabilities. To accommodate the potential sequencing errors (indels), we allowed transitions to the indel states by setting the transition probability from each match state to the insert and delete states to 0.1. In contrast, the transition probability from match to match state was 0.8. These transition probabilities were chosen *a priori* and are flexible enough to deal with indels while still preferring match-to-match transition between AAs in the proteins of the database. Remaining transition probabilities (e.g. from insertion to match, deletion to match) and all other parameters were the default values in HMMER3 (v3.3.2). To make the generated HMMs accessible to HMMER3, the HMM model fitting parameters for Viterbi, multiple ungapped segment Viterbi, and forward log-odds likelihood scores were calculated using hmmsim from HMMER3. To query the human protein database, we used PyHMMER v0.6.3 [52], a Python library binding to HMMER [51]. By default, HMMER3 checks and filters sequences with a composition bias in the AA distribution, e.g. tandem repeats, and large hydrophobic regions. However, we turned off the composition bias filter in HMMER3 for greater sensitivity [53].

### 2.7 Assessment of results

Using all proteins (N = 20,181) in our database, we simulated readouts for different cases. To identify the protein, we first constructed query HMMs based on these readouts and then scored each target protein sequence in the database. We took the protein with the highest score as the inferred protein for the given readout. For analyses using multiple fragments (readouts) per protein, we summed the HMMER scores for each target protein over all query fragments and took the highest total score to indicate the inferred protein. We used the accuracy, defined as the fraction of proteins that were correctly identified, to evaluate the performance of our method.

## 3 RESULTS

### 3.1 Effect of signal quality

We first studied the effect of signal quality on protein identification by varying the posterior of the correct AA (*p_max_*) in full-length proteins. We varied *p_max_* from 0.05, where the AAs are completely indistinguishable, to 0.9, where AAs are extremely distinguishable (Figure 2). Surprisingly, the accuracy was 0.96 (19,373 proteins correctly identified out of 20,181) even for a low *p_max_* of 0.08, and was close to 1 for *p_max_* greater than 0.09. This suggests that our method might be useful even on extremely noisy signals where there is little discrimination individual of AAs.

**Figure 2:**
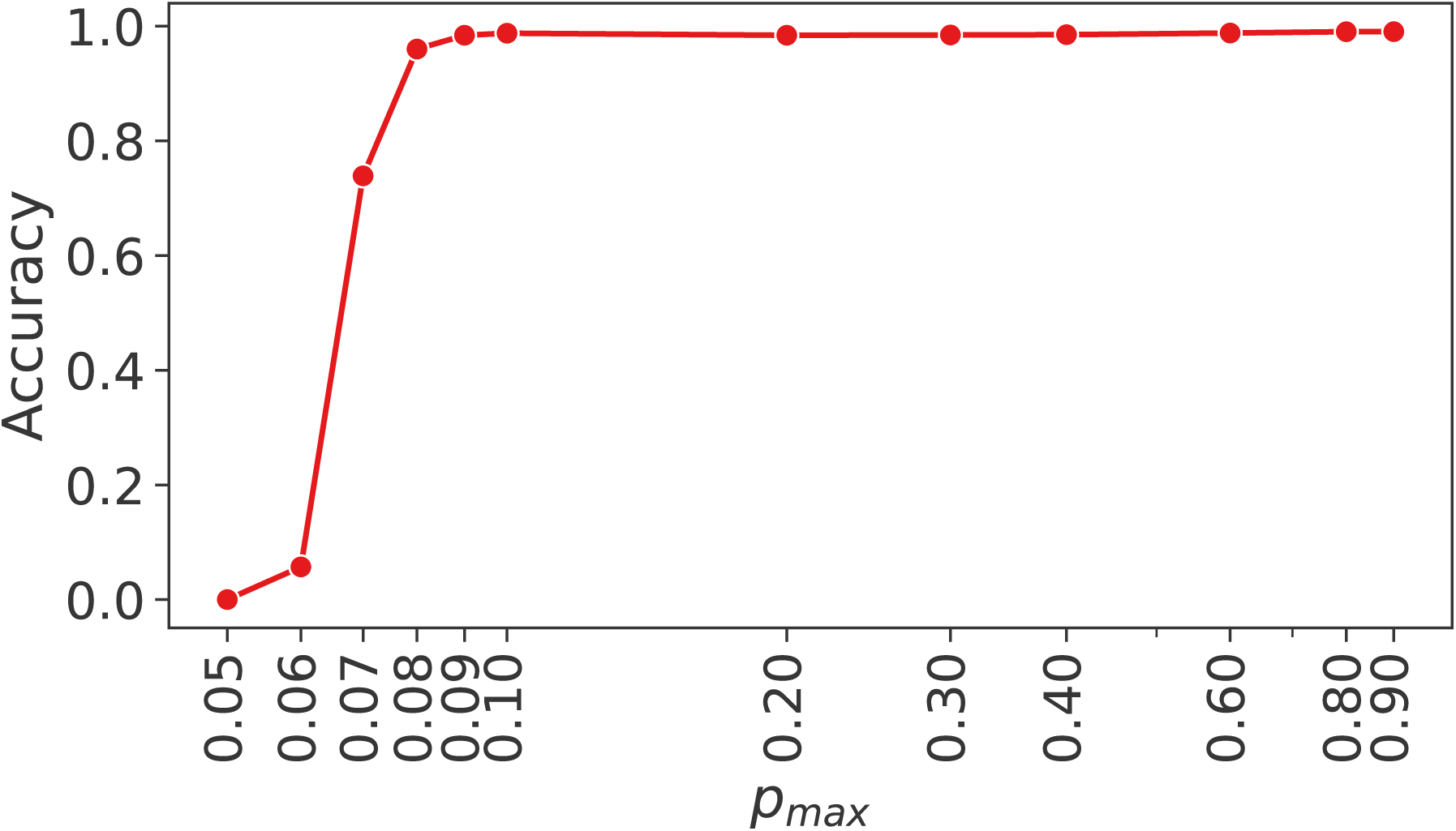
Effect of signal quality on protein identification for full-length proteins. Note the sharp increase in accuracy in the low signal quality region (*p_max_* around 0.06–0.08). The accuracy is greater than 0.95 for *p_max_*greater than 0.08. Note the logarithmic scale on the *x*-axis.

### 3.2 Effect of sequence length

To evaluate the performance on different length protein fragments, we generated random fragments of different lengths (5 AAs to 100 AAs). Since the position of a random fragment might lead to varied results, we generated 10 random fragments per sequence. In these simulations, we investigated six values for *p_max_* (0.2, 0.3, 0.4, 0.6, 0.8 and 0.9) to represent different working conditions of the sequencing device ranging from low-quality to best-quality signal outputs. The mean accuracy was greater than 0.94 for fragment lengths from 25 up to 100 AAs for all *p_max_* used (Figure 3; solid lines). For fragments of these lengths, the choice of *p_max_* had very little effect on the accuracy. This suggests that protein identification is possible with small fragments (25 AAs to 100 AAs) using devices which generate modest quality signals.

**Figure 3:**
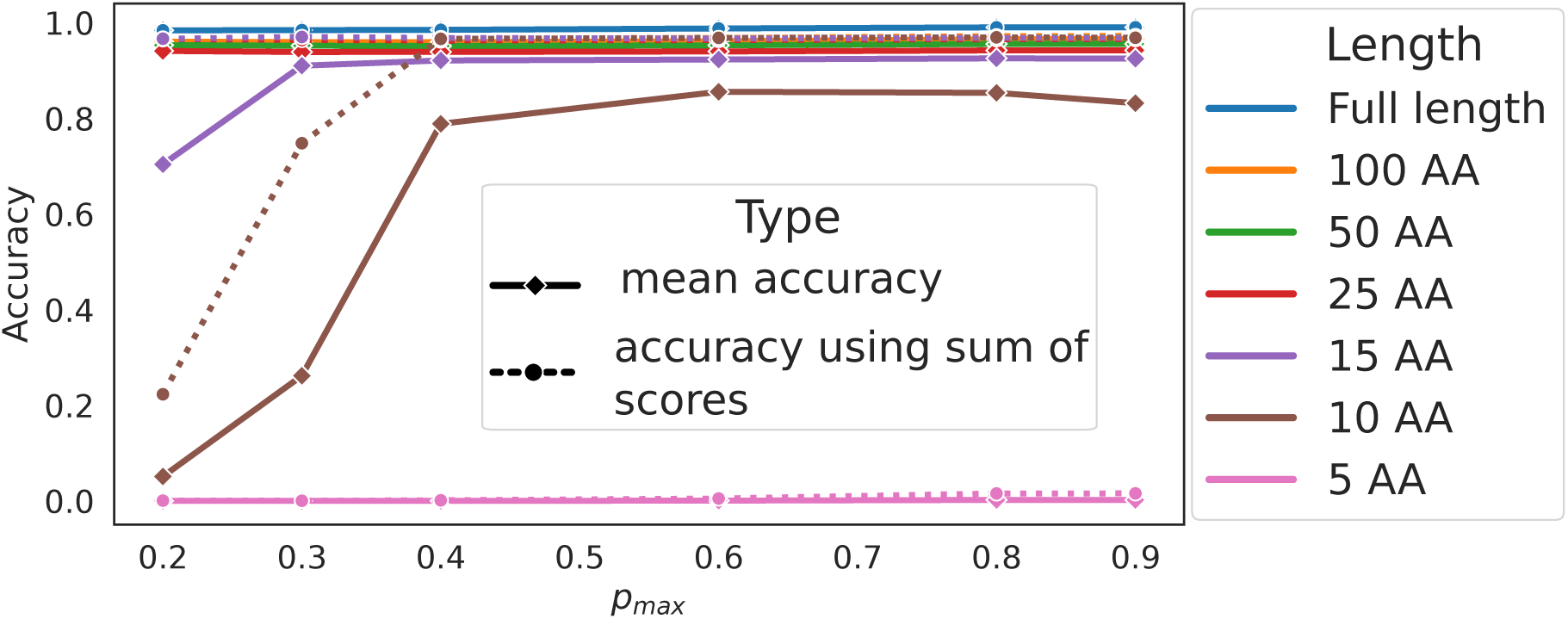
Effect of sequence length under different values of *p_max_*. The solid lines represent the mean accuracy of protein identification from single fragments, averaged over 10 random repetitions per protein. The colour of the lines indicates different lengths of the fragments. The error bars (95% CI) are smaller than the marker size. The dotted lines represent the accuracy when the sum of scores from 10 random fragments is used to infer the protein, only making a visible difference for 10- and 15-AA fragments.

On fragments shorter than 25 AAs, a higher *p_max_* was required to achieve comparable accuracy. In particular, *p_max_* above about 0.5 is needed for accuracy greater than 0.8 for fragments with as few as 10 AAs. Fragments of length 5 AAs were inadequate for protein identification, even with the highest *P_max_* values.

Additionally, in all of the above simulations, we found very little uncertainty around the mean accuracy and the error bars (95% confidence interval estimated using bootstrap, as implemented in Seaborn v0.12.2 [54]) are extremely small (Figure 3). This implies the choice of a random fragment has negligible impact on the accuracy of our method. To confirm this finding, we also assessed the proportion of these fragments that led to a correct protein identification (Supplementary Fig. S2). We find that for fragments with 25 AAs or more, irrespective of our chosen *p_max_*, the majority of sequences are identified from every one of the random fragments. This further supports our finding that each random fragment is informative, and the location bias, if any, is minimal. This also means that using a single random fragment of length 25 AAs or more in our simulations is generally sufficient for protein identification.

Next, we investigated whether the results from multiple fragments could be utilised to increase the accuracy, particularly relevant for the cases with low accuracy, i.e. with small fragment length and poorly resolved signal (< 25 AA and *p_max_* < 0.5). Using this technique, we find an increase in accuracy in almost all cases (Figure 3; dotted lines). For the longer fragments, the accuracies were already greater than 96% and improvements are not visible in the plot. Importantly, the accuracy using small fragments (e.g. 10-AA fragments) was now greater than 96% in almost all of the cases.

For 5-AA fragments, however, combining results from 10 fragments was not sufficient to identify proteins (Figure 3). We investigated whether further increasing the number of 5-AA fragments would lead to an increase in accuracy. For this, we set *p_max_* = 0.9 and increased the fragment number from 10 to 1,000. Using the sum of scores across 1,000 fragments to infer each protein, we were able to detect a modest increase in accuracy, from 0.02 to 0.10 (Supplementary Fig. S3). Thus, our method is also applicable for real-world sequencing cases where typically a sample of digested proteins would contain numerous small peptides.

### 3.3 Effect of reduced AA sets

Fluorescent labelling techniques usually label a subset of AAs to generate unique fingerprints, which can be used to identify a protein [20, 13]. Identification of a few AAs in this way, or using label-free methods with a hypothetical device that cannot observe all 20 AAs reliably [17], is relevant to new nanopore devices because it provides a way to work towards the identification of all AAs in the future while still being useful to generate useful protein signatures. Therefore, we investigated the effectiveness of our search method combined with such fingerprinting techniques. In particular, we are interested in identifying the AAs that are more informative and can provide non-ambiguous fingerprints. We first fix the working conditions of the device by setting *p_max_* to 0.8, representing a good-quality signal, or 0.2 representing a low-quality signal from the AAs. In both of these cases, we consider both the full-length and fragment (50 and 100 AAs) strategies. Based on the results already presented, these combinations provide us with settings appropriate to examine better and worse sequencing conditions, while limiting the combinatorial space.

Starting with the five most abundant AAs (L, S, E, A, and G), we gradually increase the number of these “labelled” AAs from a single AA (L) to all five (LSEAG). Surprisingly, utilising just the single AA L on full-length proteins and a good signal (*p_max_* = 0.8), our HMM search was able to identify 75% of the proteins (Figure 4A). Using additional AAs of this high-abundancy set, the accuracy was greater than 95%. This is in sharp contrast to the results from the reduced set consisting of the least abundant AAs (W, M, C, H and Y), where we required all five AAs to achieve an accuracy of 80%. We also used a third set of AAs (C, Y and K) which are often used in fluorescent labelling. C and Y are rare AAs in our database whereas K is somewhat abundant (Supplementary Fig. S1B). Compared to the previous two sets where all AAs had either high or low abundance, this set represents the labelling of AAs that both have high and low abundances. The best accuracy from this set was 80% when we used all three AAs (CYK). To simulate the practical use case where the experimenter may choose to label some of these three AAs, we ran our simulation on all possible combinations of (C, Y, K). The combinations with K had a higher performance, presumably because K is a high-frequency AA. This suggests that labelling at least one high-frequency AA might be a good strategy.

**Figure 4:**
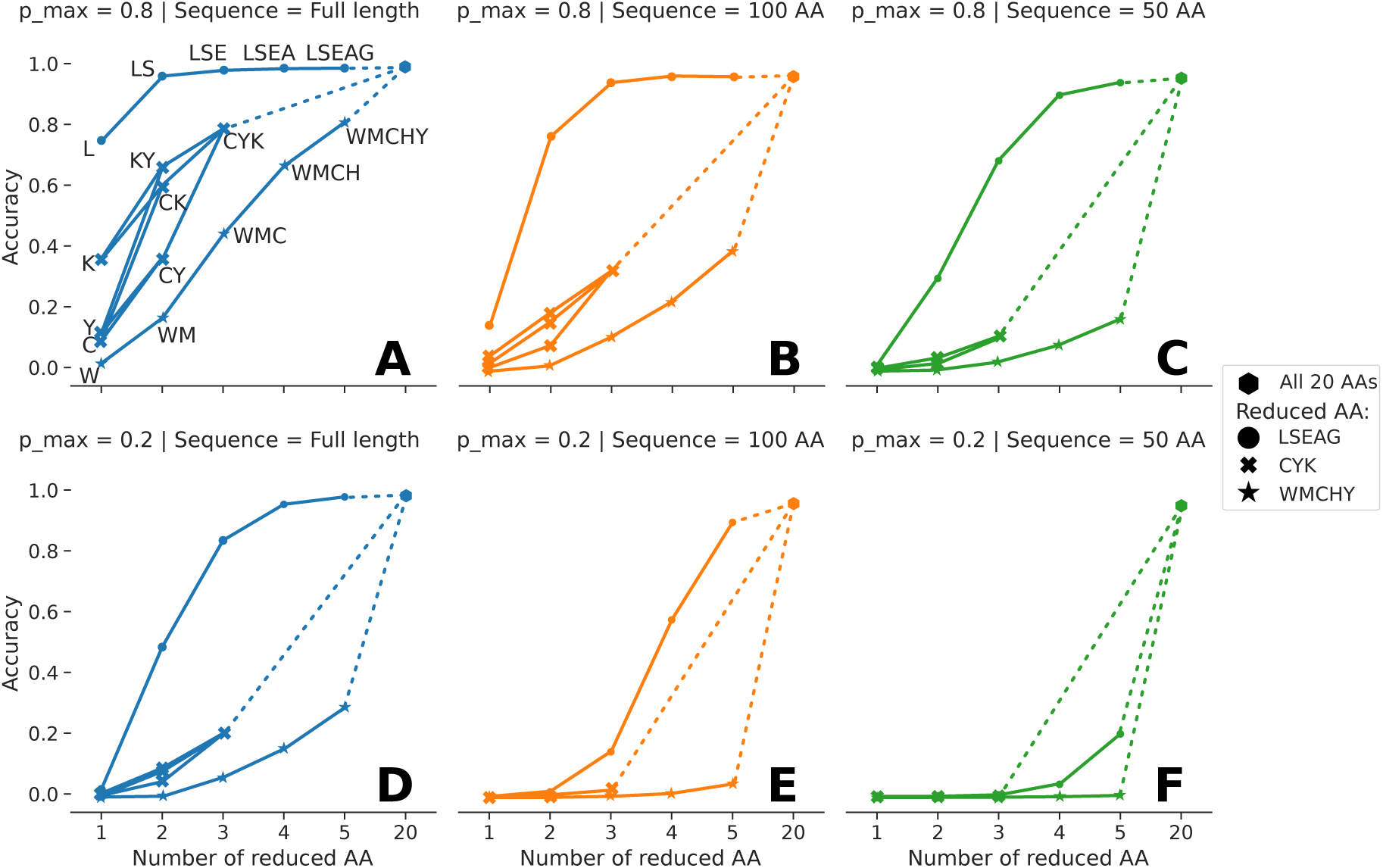
Effect of using reduced sets of AA. Three sets of reduced AA (LSEAG, CYK, WMCHY) were used with full length sequences (left column) and fragments (100 AAs and 50 AAs; centre and right, respectively), and with *p_max_* of 0.8 and 0.2 (top and bottom rows, respectively). For each AA set, the number of reduced AAs was increased from one up to all set members. The legend indicates resulting set memberships; the AAs are labelled in subfigure A and omitted in other subfigures for clarity. For comparison, the accuracy when all 20 AAs are used for each case (as in Figure 3) is also shown (hexagon marker). For the set CYK, we also used all combinations of two AA.

We repeated the above simulation for the remaining combinations of *p_max_* and sequence length (Figure 4B–F). Similar to the above result, the set LSEAG still had the highest performance followed by CYK and WMCHY. As expected, the accuracy of all three sets decreases with the decrease in sequence length (Figure 4, comparison of A–C and of D–F) and *p_max_* (compare A and D, B and E, and C and F). In particular, accuracy drops sharply for the fragments using the sets CYK and WMCHY, where we can identify less than 40% of the sequences in all the remaining cases. However, we were still able to identify 94% of the proteins using the set LSEAG on 50-AA fragments with a *p_max_* of 0.8 (Figure 4C). On further reduction of *p_max_*to 0.2, the accuracy was 0.20 (Figure 4F). In addition, this low *p_max_* was not sufficient to identify sequences from the fragments using CYK and WMCHY. These results show that given a higher quality signal, fingerprinting using more abundant AAs might still be reliable for fragments as short as 50 AAs.

### 3.4 Effect of errors

To investigate the effect of errors in protein identification, we first set the working condition of the sequencing device by fixing *p_max_* to 0.8. This simulates a good signal and makes it easier to study such effects. For worse signals, the device may be already less accurate, therefore making it harder to ascertain whether the performance is reduced due to errors or the signal quality. For the input sequence, we use full length (Figure 5), 100-(Supplementary Fig. S4) and 50-AA (Supplementary Fig. S5) fragments. For the possible AA discrimination from the device, we examine discrimination of all 20 AAs, and AAs from the reduced sets (LSEAG, WMCHY and CYK).To reduce the combinatorial space, we used all AAs from each reduced set for this analysis.

**Figure 5:**
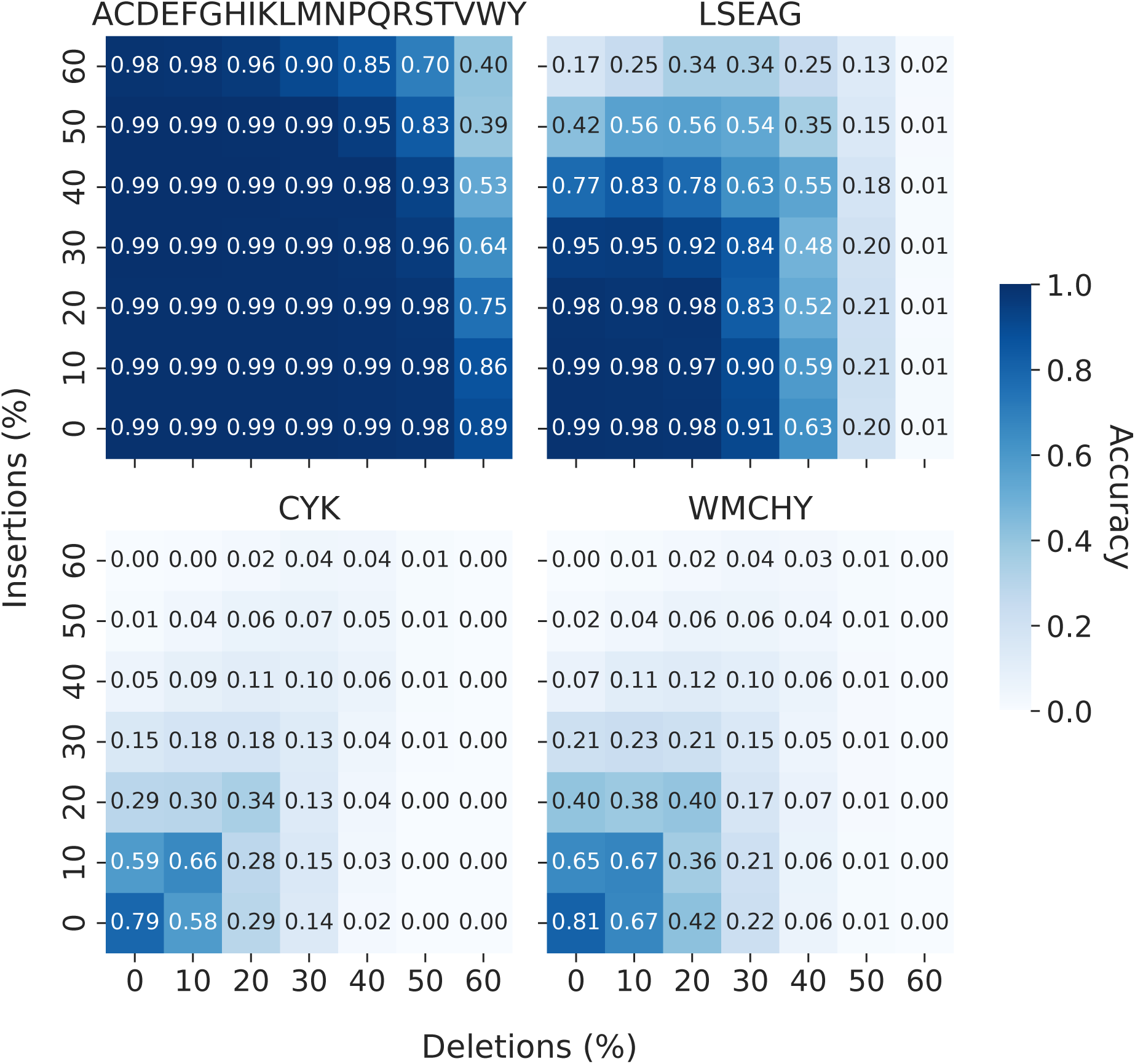
Effects of error-prone readouts on full length sequences. Heatmaps show the accuracy under different rates of insertion (*y*-axis) and deletion errors (*x*-axis) on the readouts. The subfigure titles indicate the AAs identified by the sequencer, i.e. identification of all 20 AAs (top left) and the three reduced sets of AA (remaining subfigures). For these results, we used full-length proteins from the human protein database (N=20,181). The effect of errors on protein identification using 100- and 50-AA fragments are shown in Supplementary Figs. S4, S5, respectively.

On full length sequence, with identification of all AAs, our protein identification method is extremely robust to errors (Figure 5, top left). Indel rates as high as 50% still resulted in a maximum accuracy of 0.99. However, upon further increasing the indels to 60%, the method starts to break down as reflected by the sharp decrease in accuracy (0.40). The reduced AA sets, however, are less resilient to errors. The set with the most frequent AAs (LSEAG) was the most promising and had a maximum accuracy of 0.95 at error rates of 30% (Figure 5, top right). An additional increase in error rates has a more pronounced effect on accuracy. In particular, the method starts to break down after 40% indels and the accuracy drops to nearly zero for higher error rates. In contrast, the sets with lower frequency AAs (WMCHY and CYK) had very poor tolerance to errors (Figure 5, bottom). In these sets, error rates up to 10% gave an accuracy of around 0.67 which may be considered acceptable.

On the other hand, fragments generally are less tolerant to errors and they have lower performances compared to the cases with full sequence. For example, using 100-AA fragments (Supplementary Fig. S4) with all AAs identifiable, we can identify 80 to 95% of the proteins for error rates up to 40%. With an unequal amount of insertions and deletions in readings, 60% of either of these errors still gave a maximum accuracy of 0.97. However, simultaneous increases in the insertion and deletion rates generally lead to extremely poor performance with accuracies up to around 0.2. Similar to the full length case, the high-abundance reduced set LSEAG has a better performance compared to the sets WMCHY and CYK. This set had a maximum accuracy of 0.95 for error rates of 20%. Error rates of 30% may be considered acceptable, where the maximum accuracy was around 0.63; for error rates of 50% and higher, our method was were able to identify less than 20% of the proteins. The low-abundance sets WMCHY and CYK were not useful in protein identification. They already had a very low accuracy even without any errors in the signal, and further reduction of sequence length to 50-AA fragments causes less error tolerance (Supplementary Fig. S5). For example, whereas when all AAs are identifiable the accuracies at 50% error rates can be as high as 0.95, for the reduced set LSEAG, the maximum accuracy is around 0.73 for error rates up to 20%, after which the method seems to break down.

## 4 Discussion

Protein sequencing technologies are evolving at a fast pace with many new devices and methods. To explore the potential success of these different technologies in a scenario where protein sequences are determined only with uncertainty or errors, and potentially from fragments rather than full-length proteins, we proposed a hypothetical black-box sequencing device. The readouts from our hypothetical sequencer consist of position-wise uncertainties in the AA as a distribution of posterior probabilities, with additional errors (insertions and deletions) also possible. This is comparable to position-specific probabilistic representations of protein families and domains such as profiles and motifs. Profiles, in particular, are more generalised representations because indels are allowed at every position [55, 44]. In the case of protein sequencing, indels in the signals originate from errors in the sequencing device or decoding algorithm. Therefore, HMMs are a natural way to model our probabilistic readouts.

We devised a method for protein identification that, for a given readout, builds an HMM using the posteriors as emission probabilities on a match state. Analogously to existing homology search techniques, we then query the human protein database to find the most similar sequence for a given readout. Our hypothetical device is thus suitable to study the protein identification problem by first generating data and then using it to query a database and infer the likely originating protein.

A further feature of our device is a tunable parameter, *p_max_*, corresponding to the posterior probability for the correct AA. We use this to simulate variable signal quality and decodability, irrespective of the working principles of the device. However, this is a rather simplified version of the actual operating characteristics of a real device, where signal quality could follow some complex distribution along the sequence. We also simplified indel error processes by assuming uniformly random errors along the sequence. Despite these simplifications, such a model helped us evaluate the performance of our protein identification method under diverse conditions. More complex models of sequencing device outputs could be adopted in future iterations of this research.

Successful experiments for the discrimination of all 20 AAs at a single molecule level [31] and the translocation of long peptides [26] are promising for the development of a protein sequencing device in the future that can read all AAs in full-length proteins. Simulating these conditions, we found that, in the absence of indel errors, *p_max_* as low as 0.08 — an exceedingly weak signal — was sufficient to identify more than 95% of the proteins from a single sequence (read). This suggests protein identification using our method is possible despite a marginally resolvable signal. In addition, the method can also tolerate indel rates at least as high as 50–60% if the signal quality is good (e.g. *p_max_* = 0.8).

Early devices will likely use protein fragments to mitigate problems such as nanopore clogging and limited Edman cycles. Further simplification for the device development would be to attempt the identification of a reduced set of AAs. We simulated both of these scenarios and studied the accuracy of our protein identification method. We first used random fragments of different lengths and different *p_max_*. We were able to identify at least 94% of the sequences using fragments as short as 25 AAs. Shorter fragments, containing 10 to 15 AAs, also provide a good accuracy (0.8) if the device generates good signals (*p_max_*= 0.8). Interestingly, our results also indicated that the location of a random fragment from within a protein has a negligible effect on protein identification. However, this could also be a consequence of our choice of protein database because we used canonical human proteins, where the sequences are generally unique.

Our method is easily extendable to devices which operate on multiple sequence fragments. We found that by combining scores from multiple fragments, we could significantly improve the accuracy in marginal cases.

We also studied the performance when a reduced set of identifiable AAs is used. This simulates situations such as where some AAs are fluorescent-labelled so that they are identified through the reduction of intensity after each Edman cycle, whereas the unlabeled AAs do not cause a reduction [16, 48]. Previous studies have utilised such techniques to generate protein fingerprints [19, 20, 13], although none of them proposed a general method for identifying the originating protein from a large database. In our study we investigated use of three different subsets of AAs (LSEAG, WMCHY, and CYK). Using our HMM protein identification method, the set with high-frequency AAs (LSEAG) had an extremely good performance: indeed, detection of just two AAs (LS) was sufficient to identify almost 96% of the sequences. In contrast, we need all five of the low-frequency AA set (WMCHY) to identify 80% of the sequences. The set CYK also had similar accuracy, probably because despite C and Y being rare, K is a more frequent AA in the human proteins used as our test set. For protein fragments, these sets gave a lower performance depending on *p_max_* and the length; however, the set LSEAG still had the highest accuracy. Thus, we conclude that if labelling is to be used, labelling more frequent AAs would be advantageous in our method for protein identification and could give excellent results.

We further investigated the impact of possible errors from the sequencing device by generating indel-prone readings. We found that on devices where there is discrimination of all 20 AAs, even with indel rates as high as 40% to 50% our method is successful irrespective of whether full length or fragments were used. In many cases studied, accuracy was greater than 0.90. However, reduced AA sets were not as robust to errors. The set (LSEAG) with highly abundant AAs was the most promising and had acceptable performances for error rates up to 20–30% depending upon the length of the input sequence. Therefore, our method can be used on devices with low to moderate error rates.

In conclusion, we used a hypothetical sequencing device to explore the ability of a novel, HMM-based method to identify proteins under different types of possible readouts. In many of the cases, we had good accuracy, which suggests that our method could be successfully applied to different sequencing devices in the future. Our results could also help develop new strategies for protein sequencing devices.

## 5 Data availability

The code and scripts to reproduce our results are available at https://github.com/goldman-gp-ebi/ protein-identification-manuscript.

## 7 Author contributions

N.G. conceived the study and method, and supervised the study. B.K.B implemented the method and performed all analyses. N.G and B.K.B drafted, reviewed, edited, and approved the manuscript.

## References

[1] Erwin L van Dijk, Hélène Auger, Yan Jaszczyszyn, and Claude Thermes. Ten years of next-generation sequencing technology. Trends Genet., 30(9):418–426, September 2014.

[2] Anthony Rhoads and Kin Fai Au. PacBio sequencing and its applications. Genomics, Proteomics & Bioinformatics, 13(5):278–289, October 2015.

[3] James M Heather and Benjamin Chain. The sequence of sequencers: the history of sequencing DNA. Genomics, 107(1):1–8, January 2016.

[4] Miten Jain, Hugh E Olsen, Benedict Paten, and Mark Akeson. The Oxford Nanopore MinION: delivery of nanopore sequencing to the genomics community. Genome Biol., 17(1):239, November 2016.

[5] Yunhao Wang, Yue Zhao, Audrey Bollas, Yuru Wang, and Kin Fai Au. Nanopore sequencing technology, bioinformatics and applications. Nat. Biotechnol., 39(11):1348–1365, November 2021.

[6] Yusuke Goto, Rena Akahori, Itaru Yanagi, and Ken-Ichi Takeda. Solid-state nanopores towards single-molecule DNA sequencing. J. Hum. Genet., 65(1):69–77, January 2020.

[7] Kristoffer Sahlin and Paul Medvedev. Error correction enables use of Oxford Nanopore technology for reference-free transcriptome analysis. Nat. Commun., 12(1):2, January 2021.

[8] Franka J Rang, Wigard P Kloosterman, and Jeroen de Ridder. From squiggle to basepair: computational approaches for improving nanopore sequencing read accuracy. Genome Biol., 19(1):90, July 2018.

[9] Henry Brinkerhoff, Albert S W Kang, Jingqian Liu, Aleksei Aksimentiev, and Cees Dekker. Multiple rereads of single proteins at single-amino acid resolution using nanopores. Science, 374(6574):1509–1513, December 2021.

[10] Brian D Reed, Michael J Meyer, Valentin Abramzon, Omer Ad, Omer Ad, Pat Adcock, Faisal R Ahmad, Gün Alppay, James A Ball, James Beach, Dominique Belhachemi, Anthony Bellofiore, Michael Bellos, Juan Felipe Beltrán, Andrew Betts, Mohammad Wadud Bhuiya, Kristin Blacklock, Robert Boer, David Boisvert, Norman D Brault, Aaron Buxbaum, Steve Caprio, Changhoon Choi, Thomas D Christian, Robert Clancy, Joseph Clark, Thomas Connolly, Kathren Fink Croce, Richard Cullen, Mel Davey, Jack Davidson, Mohamed M Elshenawy, Michael Ferrigno, Daniel Frier, Saketh Gudipati, Stephanie Hamill, Zhaoyu He, Sharath Hosali, Haidong Huang, Le Huang, Ali Kabiri, Gennadiy Kriger, Brittany Lathrop, An Li, Peter Lim, Stephen Liu, Feixiang Luo, Caixia Lv, Xiaoxiao Ma, Evan McCormack, Michele Millham, Roger Nani, Manjula Pandey, John Parillo, Gayatri Patel, Douglas H Pike, Kyle Preston, Adeline Pichard-Kostuch, Kyle Rearick, Todd Rearick, Marco Ribezzi-Crivellari, Gerard Schmid, Jonathan Schultz, Xinghua Shi, Badri Singh, Nikita Srivastava, Shannon F Stewman, T R Thurston, T R Thurston, Philip Trioli, Jennifer Tullman, Xin Wang, Yen-Chih Wang, Eric A G Webster, Zhizhuo Zhang, Jorge Zuniga, Smita S Patel, Andrew D Griffiths, Antoine M van Oijen, Michael McKenna, Matthew D Dyer, and Jonathan M Rothberg. Real-time dynamic single-molecule protein sequencing on an integrated semiconductor device. Science, 378(6616):186–192, October 2022.

[11] Zheng-Li Hu, Ming-Zhu Huo, Yi-Lun Ying, and Yi-Tao Long. Biological nanopore approach for single-molecule protein sequencing. Angew. Chem. Int. Ed., 60(27):14738–14749, June 2021.

[12] Shengli Zhang, Gang Huang, Roderick Corstiaan Abraham Versloot, Bart Marlon Herwig Bruininks, Paulo Cesar Telles de Souza, Siewert-Jan Marrink, and Giovanni Maglia. Bottom-up fabrication of a proteasome-nanopore that unravels and processes single proteins. Nat. Chem., 13(12):1192–1199, December 2021.

[13] Jagannath Swaminathan, Alexander A Boulgakov, Erik T Hernandez, Angela M Bardo, James L Bachman, Joseph Marotta, Amber M Johnson, Eric V Anslyn, and Edward M Marcotte. Highly parallel single-molecule identification of proteins in zeptomole-scale mixtures. Nat. Biotechnol., 36(11):1076–1082, October 2018.

[14] Brendan M Floyd and Edward M Marcotte. Protein sequencing, one molecule at a time. Annu. Rev. Biophys., 51:181–200, May 2022.

[15] Keisuke Motone and Jeff Nivala. Not if but when nanopore protein sequencing meets single-cell proteomics. Nat. Methods, 20(3):336–338, March 2023.

[16] Laura Restrepo-Pérez, Chirlmin Joo, and Cees Dekker. Paving the way to single-molecule protein sequencing. Nat. Nanotechnol., 13(9):786–796, September 2018.

[17] Yingqi Zhao, Marzia Iarossi, Angela Federica De Fazio, Jian-An Huang, and Francesco De Angelis. Label-free optical analysis of biomolecules in solid-state nanopores: toward single-molecule protein sequencing. ACS Photonics, 9(3):730–742, March 2022.

[18] Katalin F Medzihradszky and Robert J Chalkley. Lessons in *de novo* peptide sequencing by tandem mass spectrometry. Mass Spectrometry Reviews, 34(1):43–63, January/February 2015.

[19] Jagannath Swaminathan, Alexander A Boulgakov, and Edward M Marcotte. A theoretical justification for single molecule peptide sequencing. PLoS Comput. Biol., 11(2):e1004080, February 2015.

[20] Yao Yao, Margreet Docter, Jetty van Ginkel, Dick de Ridder, and Chirlmin Joo. Single-molecule protein sequencing through fingerprinting: computational assessment. Phys. Biol., 12(5):055003, August 2015.

[21] Jetty van Ginkel, Mike Filius, Malwina Szczepaniak, Pawel Tulinski, Anne S Meyer, and Chirlmin Joo. Single-molecule peptide fingerprinting. Proc. Natl. Acad. Sci. U. S. A., 115(13):3338–3343, March 2018.

[22] Erik T Hernandez, Jagannath Swaminathan, Edward M Marcotte, and Eric V Anslyn. Solution-phase and solid-phase sequential, selective modification of side chains in KDYWEC and KDYWE as models for usage in single-molecule protein sequencing. New J. Chem., 41(2):462–469, January 2017.

[23] Jeff Nivala, Douglas B Marks, and Mark Akeson. Unfoldase-mediated protein translocation through an *α*-hemolysin nanopore. Nat. Biotechnol., 31(3):247–250, March 2013.

[24] Chan Cao, Nuria Cirauqui, Maria Jose Marcaida, Elena Buglakova, Alice Duperrex, Aleksandra Radenovic, and Matteo Dal Peraro. Single-molecule sensing of peptides and nucleic acids by engineered aerolysin nanopores. Nat. Commun., 10(1):4918, October 2019.

[25] Mazdak Afshar Bakshloo, John J Kasianowicz, Manuela Pastoriza-Gallego, Jérôme Mathé, Régis Daniel, Fabien Piguet, and Abdelghani Oukhaled. Nanopore-based protein identification. Journal of the American Chemical Society, 144(6):2716–2725, February 2022.

[26] Luning Yu, Xinqi Kang, Fanjun Li, Behzad Mehrafrooz, Amr Makhamreh, Ali Fallahi, Joshua C Foster, Aleksei Aksimentiev, Min Chen, and Meni Wanunu. Unidirectional single-file transport of full-length proteins through a nanopore. Nat. Biotechnol., 41(8):1130–1139, January 2023.

[27] Hadjer Ouldali, Kumar Sarthak, Tobias Ensslen, Fabien Piguet, Philippe Manivet, Juan Pelta, Jan C Behrends, Aleksei Aksimentiev, and Abdelghani Oukhaled. Electrical recognition of the twenty proteinogenic amino acids using an aerolysin nanopore. Nat. Biotechnol., 38(2):176–181, February 2020.

[28] Laura Restrepo-Pérez, Shalini John, Aleksei Aksimentiev, Chirlmin Joo, and Cees Dekker. SDS-assisted protein transport through solid-state nanopores. Nanoscale, 9(32):11685–11693, July 2017.

[29] Wang Li, Juan Zhou, Nicolò Maccaferri, Roman Krahne, Kang Wang, and Denis Garoli. Enhanced optical spectroscopy for multiplexed DNA and protein-sequencing with plasmonic nanopores: challenges and prospects. Anal. Chem., 94(2):503–514, January 2022.

[30] Xiaowen Liu, Zhuxin Dong, and Gregory Timp. Calling the amino acid sequence of a protein/peptide from the nanospectrum produced by a sub-nanometer diameter pore. Sci. Rep., 12(1):17853, October 2022.

[31] Jian-An Huang, Mansoureh Z Mousavi, Giorgia Giovannini, Yingqi Zhao, Aliaksandr Hubarevich, Miguel A Soler, Walter Rocchia, Denis Garoli, and Francesco De Angelis. Multiplexed discrimination of single amino acid residues in polypeptides in a single SERS hot spot. Angew. Chem. Int. Ed., 59(28):11423–11431, July 2020.

[32] Judith Langer, Dorleta Jimenez de Aberasturi, Javier Aizpurua, Ramon A Alvarez-Puebla, Bap-tiste Auguié, Jeremy J Baumberg, Guillermo C Bazan, Steven E J Bell, Anja Boisen, Alexandre G Brolo, Jaebum Choo, Dana Cialla-May, Volker Deckert, Laura Fabris, Karen Faulds, F Javier García de Abajo, Royston Goodacre, Duncan Graham, Amanda J Haes, Christy L Haynes, Christian Huck, Tamitake Itoh, Mikael Käll, Janina Kneipp, Nicholas A Kotov, Hua Kuang, Eric C Le Ru, Hiang Kwee Lee, Jian-Feng Li, Xing Yi Ling, Stefan A Maier, Thomas Mayerhöfer, Martin Moskovits, Kei Murakoshi, Jwa-Min Nam, Shuming Nie, Yukihiro Ozaki, Isabel Pastoriza-Santos, Jorge Perez-Juste, Juergen Popp, Annemarie Pucci, Stephanie Reich, Bin Ren, George C Schatz, Timur Shegai, Sebastian Schlücker, Li-Lin Tay, K George Thomas, Zhong-Qun Tian, Richard P Van Duyne, Tuan Vo-Dinh, Yue Wang, Katherine A Willets, Chuanlai Xu, Hongxing Xu, Yikai Xu, Yuko S Yamamoto, Bing Zhao, and Luis M Liz-Marzán. Present and future of surface-enhanced Raman scattering. ACS Nano, 14(1):28–117, January 2020.

[33] Juan Zhou, Qing Lan, Wang Li, Li-Na Ji, Kang Wang, and Xing-Hua Xia. Single molecule protein segments sequencing by a plasmonic nanopore. Nano Letters, 23(7):2800–2807, March 2023.

[34] Nicholas Callahan, Jennifer Tullman, Zvi Kelman, and John Marino. Strategies for development of a next-generation protein sequencing platform. Trends Biochem. Sci., 45(1):76–89, January 2020.

[35] UniProt Consortium. UniProt: the universal protein knowledgebase in 2021. Nucleic Acids Res., 49(D1):D480–D489, January 2021.

[36] Jacob Schreiber and Kevin Karplus. Analysis of nanopore data using hidden Markov models. Bioinformatics, 31(12):1897–1903, June 2015.

[37] Vladimí Bža, Broňa Brejová, and Tomáš Vinař. DeepNano: deep recurrent neural networks for base calling in MinION nanopore reads. PLoS One, 12(6):e0178751, June 2017.

[38] Matei David, L J Dursi, Delia Yao, Paul C Boutros, and Jared T Simpson. Nanocall: an open source basecaller for Oxford Nanopore sequencing data. Bioinformatics, 33(1):49–55, January 2017.

[39] Haotian Teng, Minh Duc Cao, Michael B Hall, Tania Duarte, Sheng Wang, and Lachlan J M Coin. Chiron: translating nanopore raw signal directly into nucleotide sequence using deep learning. Gigascience, 7(5):giy037, May 2018.

[40] Ryan R Wick, Louise M Judd, and Kathryn E Holt. Performance of neural network basecalling tools for Oxford Nanopore sequencing. Genome Biol., 20(1):129, June 2019.

[41] Yuwei Bao, Jack Wadden, John R Erb-Downward, Piyush Ranjan, Weichen Zhou, Torrin L McDonald, Ryan E Mills, Alan P Boyle, Robert P Dickson, David Blaauw, and Joshua D Welch. SquiggleNet: real-time, direct classification of nanopore signals. Genome Biol., 22(1):298, October 2021.

[42] Don Neumann, Anireddy S N Reddy, and Asa Ben-Hur. RODAN: a fully convolutional architecture for basecalling nanopore RNA sequencing data. BMC Bioinformatics, 23(1):142, April 2022.

[43] Anjana Senanayake, Hasindu Gamaarachchi, Damayanthi Herath, and Roshan Ragel. DeepSelectNet: deep neural network based selective sequencing for oxford nanopore sequencing. BMC Bioinformatics, 24(1):31, January 2023.

[44] S R Eddy. Profile hidden Markov models. Bioinformatics, 14(9):755–763, January 1998.

[45] Jeff Nivala, Logan Mulroney, Gabriel Li, Jacob Schreiber, and Mark Akeson. Discrimination among protein variants using an unfoldase-coupled nanopore. ACS Nano, 8(12):12365–12375, December 2014.

[46] Nuno Bandeira, Victoria Pham, Pavel Pevzner, David Arnott, and Jennie R Lill. Automated *de novo* protein sequencing of monoclonal antibodies. Nat. Biotechnol., 26(12):1336–1338, December 2008.

[47] Giovanni Di Muccio, Aldo Eugenio Rossini, Daniele Di Marino, Giuseppe Zollo, and Mauro Chinappi. Insights into protein sequencing with an *α*-Hemolysin nanopore by atomistic simulations. Sci. Rep., 9(1):6440, April 2019.

[48] Haihan He, Chuhong Wu, Muhammad Saqib, and Rui Hao. Single-molecule fluorescence methods for protein biomarker analysis. Anal. Bioanal. Chem., 415(18):3655–3669, July 2023.

[49] Haowen Zhang, Haoran Li, Chirag Jain, Haoyu Cheng, Kin Fai Au, Heng Li, and Srinivas Aluru. Real-time mapping of nanopore raw signals. Bioinformatics, 37(Suppl 1):i477–i483, July 2021.

[50] Richard Durbin, Sean R Eddy, Anders Krogh, and Graeme Mitchison. Biological sequence analysis: Probabilistic models of proteins and nucleic acids. Cambridge University Press, Cambridge, England, 1998.

[51] HMMER. http://hmmer.org. Accessed: 2022-10-19.

[52] Martin Larralde and Georg Zeller. PyHMMER: a python library binding to HMMER for efficient sequence analysis. Bioinformatics, 39(5):btad214, May 2023.

[53] HMMER user guide. http://eddylab.org/software/hmmer/Userguide.pdf. Accessed: 2023-9-11.

[54] Michael L. Waskom. seaborn: statistical data visualization. Journal of Open Source Software, 6(60):3021, April 2021.

[55] M Gribskov, A D McLachlan, and D Eisenberg. Profile analysis: detection of distantly related proteins. Proc. Natl. Acad. Sci. U. S. A., 84(13):4355–4358, July 1987.

