## Supplementary Figures for "A generalised protein identification method for novel and diverse sequencing technologies"

**Supplementary materials**

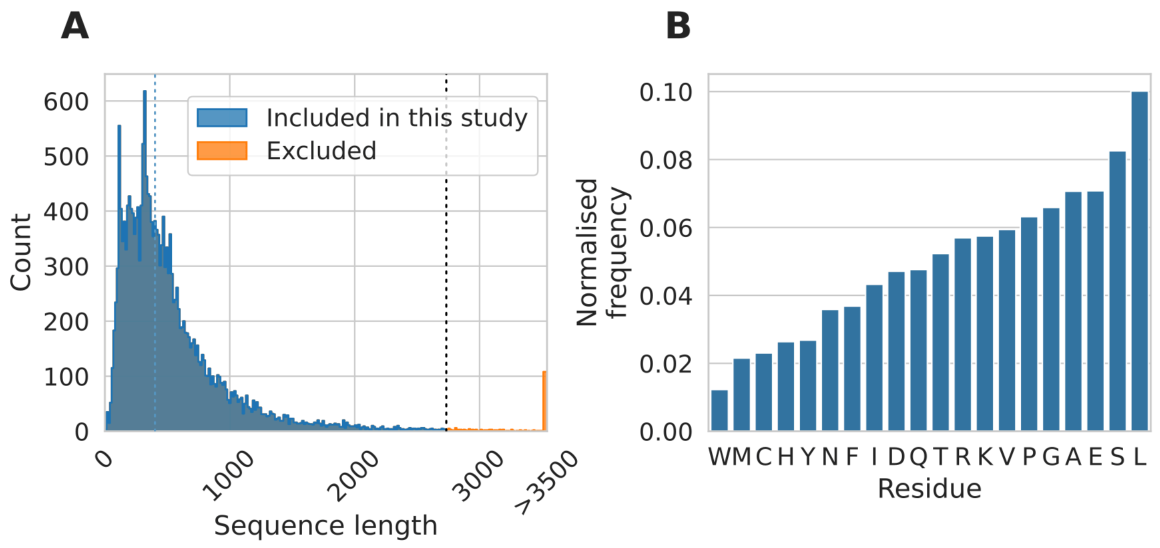


**Fig. S1. Summary statistics for the sequences from the UniProt human protein database. A)** Protein sequence length distribution. We excluded 204 sequences (orange) with length greater than the 99th percentile (denoted by vertical black dotted line) and retained the remaining sequences (blue, N=20,181) in our study. The median length of the remaining sequences is 411 residues (vertical blue dotted line). **B)** Frequency of residues in the sequences used in this study (total of 10,533,212 residues).

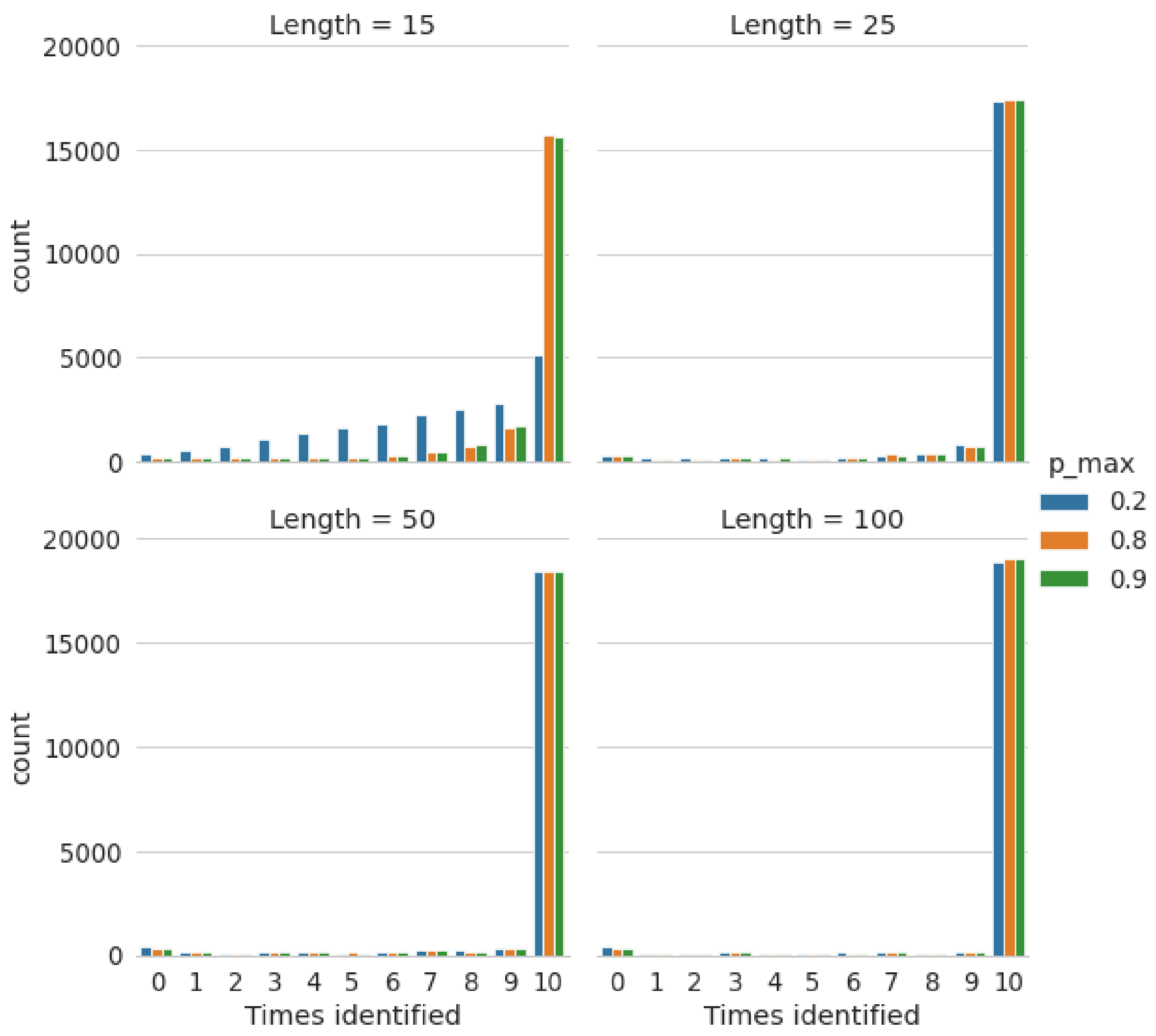


**Fig. S2. Distributions of numbers of times that human proteome database sequences (N=20,181) were identified from 10 random fragments.** For fragments of length 25 AA and above, irrespective of $p_{max}$, the majority of the protein sequences were identified from every one of the 10 random fragments.

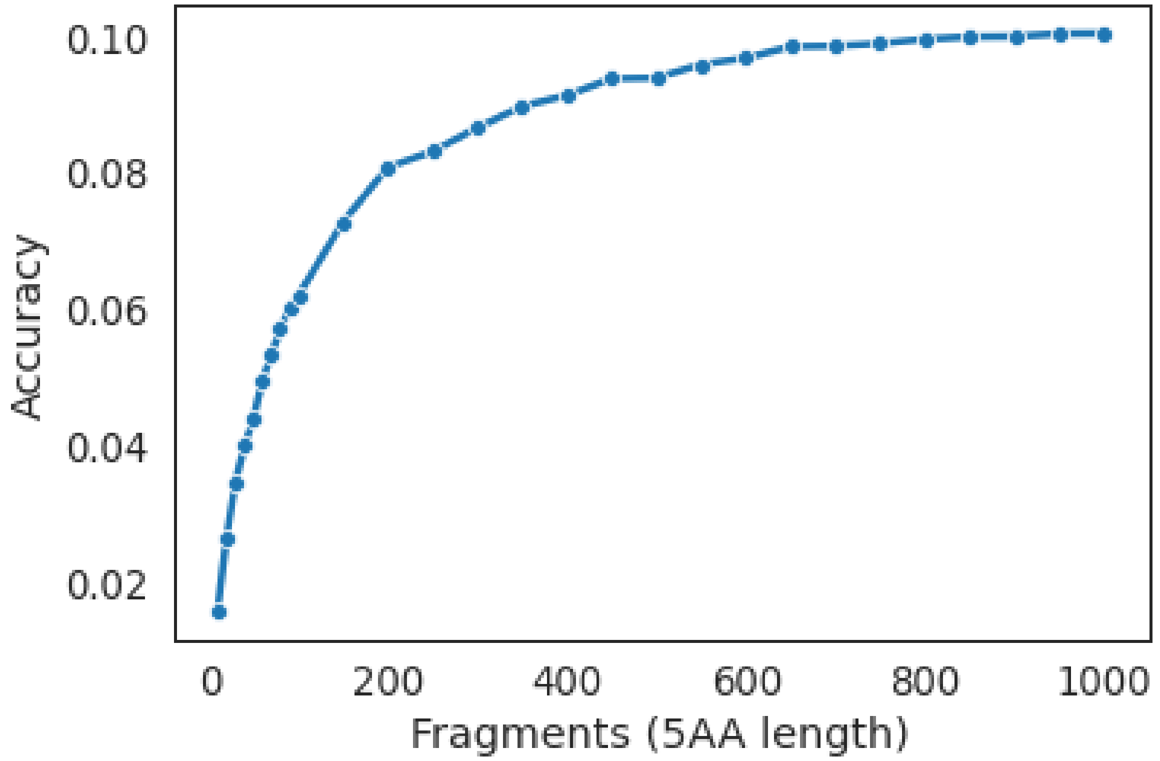


**Fig. S3. Low protein identification accuracy can be increased with increase in number of fragments.** For this result, we used $p_{max}$ = 0.9, and increased the number of 5-AA fragments per sequence from 10 to 1,000.

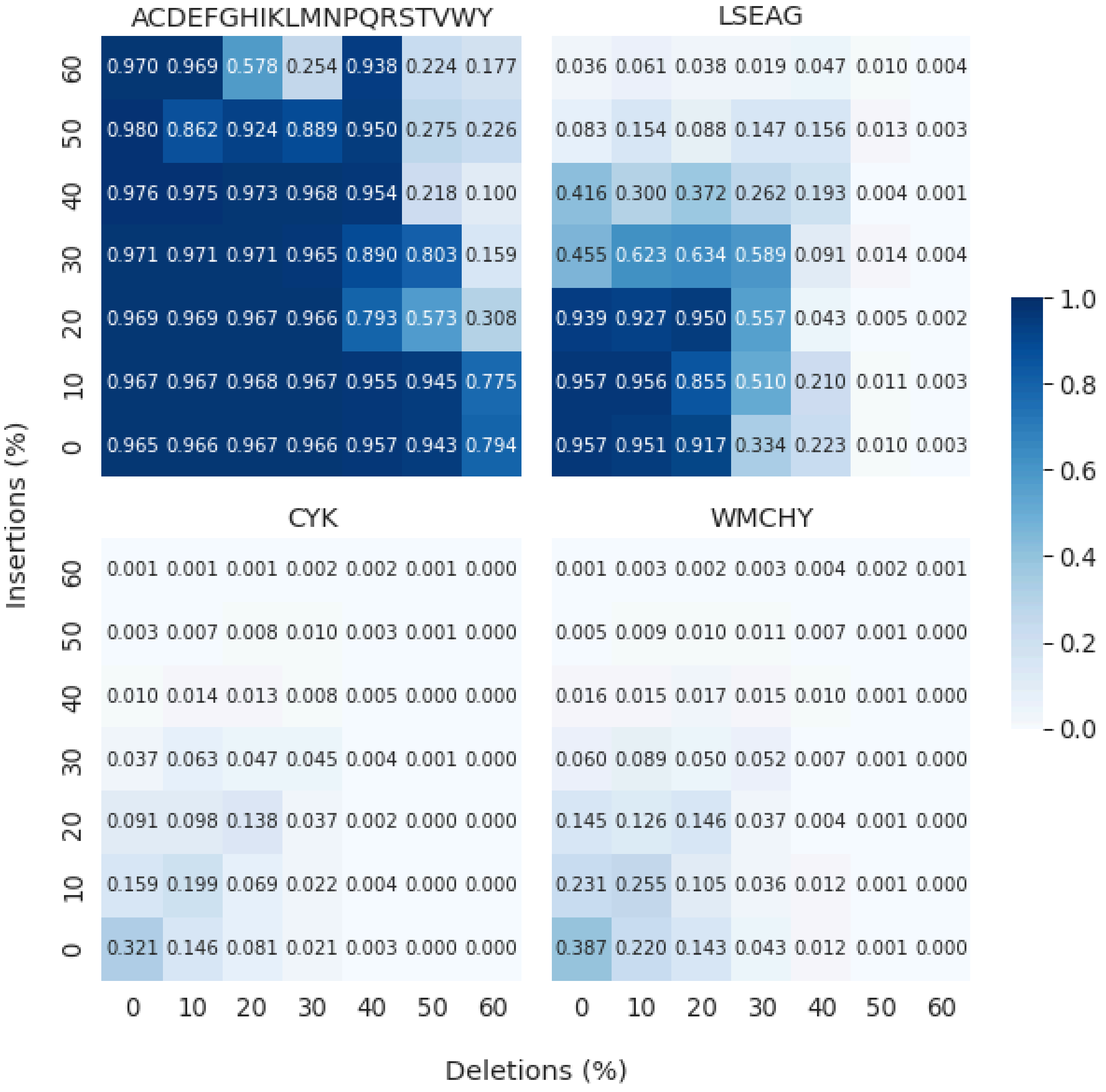


**Fig. S4. Effects of error-prone readouts on 100 AA fragments.** Heatmaps show the accuracy under different levels of insertion (*y*-axis) and deletion errors (*x*-axis) on the readouts. The subfigure titles indicate the AAs identified by the sequencer, i.e. identification of all AAs (top left) and the three reduced sets of AA (remaining subfigures). For these results, we used protein fragments with length 100 AAs from the human protein database (N=20,181).

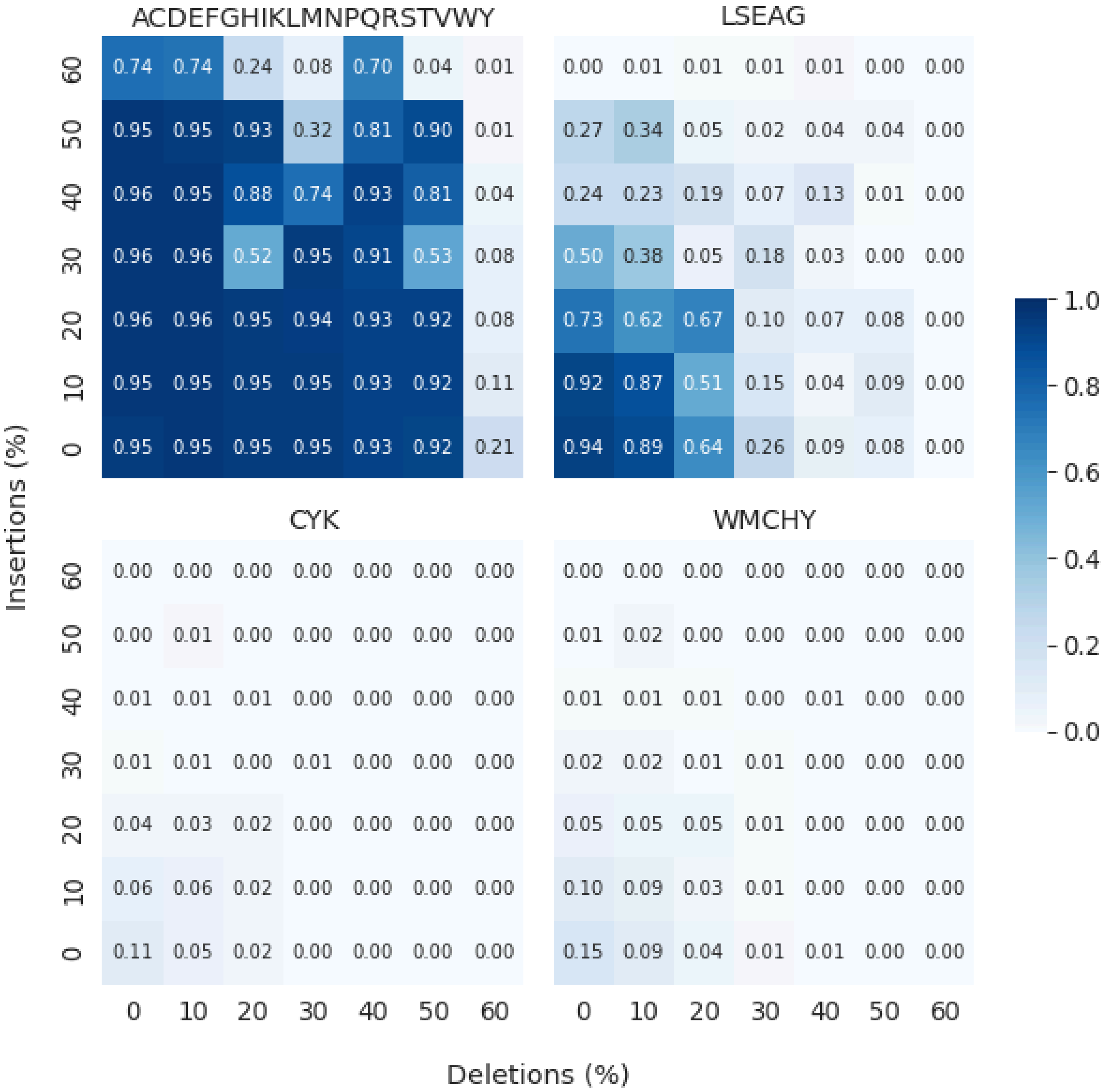


**Fig. S5. Effects of error-prone readouts on 50 AA fragments.** Heatmaps show the accuracy under different levels of insertion (*y*-axis) and deletion errors (*x*-axis) on the readouts. The subfigure titles indicate the AAs identified by the sequencer, i.e. identification of all AAs (top left) and the three reduced sets of AA (remaining subfigures). For these results, we used protein fragments with length 50 AAs from the human protein database (N=20,181).
